# Megaplasmids on the Rise: Combining Sequencing Approaches to Fully Resolve a Carbapenemase-Encoding Plasmid in a Proposed Novel *Pseudomonas* Species

**DOI:** 10.1101/601898

**Authors:** João Botelho, Cédric Lood, Sally R. Partridge, Vera van Noort, Rob Lavigne, Filipa Grosso, Luísa Peixe

## Abstract

Horizontal transfer of plasmids plays a pivotal role in the dissemination of antibiotic resistance genes and emergence of multidrug-resistant bacteria. Sequencing of plasmids is thus paramount for the success of accurate epidemiological tracking strategies in the hospital setting and routine surveillance. Here, we combine Nanopore and Illumina sequencing to fully assemble a carbapenemase-encoding megaplasmid carried by a clinical isolate belonging to a putative novel *Pseudomonas* species. FFUP_PS_41 has a multidrug resistance phenotype and was initially identified as *Pseudomonas putida*, but an average nucleotide identity below the cut-off for species delineation suggests a new species related to the *P. putida* phylogenetic group. FFUP_PS_41 harbors a 498,516-bp untypable megaplasmid (pJBCL41) with low similarity compared with publicly available plasmids. pJBCL41 contains a full set of genes for self-transmission and genes predicted to be responsible for plasmid replication, partitioning, maintenance and heavy metal resistance. pJBCL41 carries a class 1 integron with the |*aacA7*|*bla*_VIM-2_|*aacA4*| cassette array (In103) located within a defective Tn*402*-like transposon that forms part of a 50,273-bp mosaic region bound by 38-bp inverted repeats typical of the Tn*3* family and flanked by 5-bp direct repeats. This region is composed of different elements, including additional transposon fragments, five insertion sequences and a Tn*3*-Derived Inverted-Repeat Miniature Element. The hybrid Nanopore/Illumina approach resulted in contiguous assemblies and allowed us to fully resolve a carbapenemase-encoding megaplasmid from *Pseudomonas* spp. The identification of novel megaplasmids will shed a new light on the evolutionary effects of gene transfer and the selective forces driving AR.

## Introduction

Bacteria can become resistant to antibiotics through chromosomal mutations and by the acquisition of resistance genes carried on mobile genetic elements, including plasmids and integrative and conjugative elements [1]. Plasmids are autonomous self-replicating elements that drive the horizontal transfer (HGT) of antibiotic resistance genes from cell to cell by conjugation [2–5]. The mobility of a plasmid depends on the set of genes that it carries, and these extrachromosomal elements may be conjugative, mobilizable or non-transmissible [2, 3]. Conjugative plasmids carry all the machinery necessary for self-propagation: i) a relaxase, a key protein in conjugation; ii) an origin of transfer (*oriT*); iii) a set of genes encoding for the type-IV secretion system (T4SS); and iv) a gene encoding a type-IV coupling protein (T4CP) [2, 3]. Mobilizable plasmids lack the complete set of genes encoding the T4SS and may use the conjugative apparatus of a helper plasmid present in the cell to be successfully transferred. Conjugative plasmids tend to be low copy number and large, whereas mobilizable plasmids are frequently high copy number and smaller (<30 kb) [2, 3]. The term megaplasmids [6] has been used for very large replicons (>350 kb) which, in contrast to chromids [7], do not carry essential core genes. Megaplasmids frequently have mosaic structures, carrying genetic modules that originate from different ancestral sources [8]. The formation of mosaic plasmids may be influenced by several factors, such as the abundance of conjugative plasmids and transposons, selection pressures, incompatibility groups and the host’s tolerance of foreign DNA. According to the plasmid hypothesis, megaplasmids are the evolutionary precursors of chromids, due to the amelioration of genomic signatures to those of the chromosomal partner and the acquisition of essential genes [7].

To date, fourteen incompatibility groups (IncP-1 to IncP-14) have been characterized amongst *Pseudomonas* plasmids [9, 10]. Narrow host range plasmids comprise IncP types -2, -5, -7, -10, -12 and -13 and cannot be transferred into *Escherichia coli*. In contrast, other groups display a broad host range, as they are also included in the typing scheme for *Enterobacteriaceae* plasmids: IncP-1 (IncP), IncP-4 (IncQ) and IncP-6 (IncG) [9, 10]. Unlike *Enterobacteriaceae* plasmids, no replicon-based PCR typing of *Pseudomonas* plasmids has been created yet.

Plasmids may harbor accessory module(s) that provide adaptive advantage(s) for their host, such as virulence-encoding factors and antibiotic resistance genes [9, 11–13]. Sequencing of plasmids is thus paramount to the success of accurate epidemiological tracking strategy in the hospital setting and routine surveillance, helping to identify transmission routes and to prevent future outbreaks [14–19]. The advent of WGS has enabled the *in silico* analysis of a wide array of plasmids, most of them from assembly of short-read sequencing data [20–24]. However, fully resolving plasmids with short-read sequencing technologies remains challenging due to the presence of numerous long repeated regions [25], and currently the most accurate approach to assemble these plasmids is to use a combination of short-read and long-read methods [14–19, 26, 27]. Here, we combined Nanopore and Illumina sequencing to fully assemble a carbapenemase-encoding megaplasmid carried by an isolate belonging to a putative novel *Pseudomonas* species.

## Material and Methods

### Bacterial Isolate

Isolate FFUP_PS_41 was obtained in 2008 from endotracheal tube secretions of a patient with pneumonia admitted to the Neonatal/Pediatric Intensive Care unit of Centro Hospitalar do Porto – Hospital de Santo António, in Porto, Portugal, as part of regular surveillance of carbapenemase-producers among clinical isolates.

FFUP_PS_41 was initially identified as *Pseudomonas putida* by VITEK-2 (bioMérieux) and later re-classified by pair-wise average nucleotide identity based on BLAST+ (ANIb) using JSpeciesWS v3.0.20 and PyANI v0.2.7 (https://github.com/widdowquinn/pyani) [28–30]. Antimicrobial susceptibility testing was conducted by standard disc diffusion and broth microdilution (for colistin) methods, according to EUCAST guidelines (http://www.eucast.org/).

### Whole-Plasmid Sequencing and Bioinformatics

Genomic DNA from FFUP_PS_41 was extracted using a QIAamp DNA Mini Kit (Qiagen, Hilden, Germany) according to the manufacturer’s instructions. Sequencing libraries were prepared using Illumina Nextera and the 1D ligation library approach from Oxford Nanopore Technology (ONT). Libraries were sequenced on the Illumina HiSeq 2500 sequencer or the MinION sequencer from ONT equipped with a flowcell of chemistry type R9.4, respectively.

Illumina reads were verified for quality using FastQC and Trimmomatic [31, 32], while MinION reads were processed with ONT’s albacore v2.3.0 followed by demultiplexing using porechop v0.2.3. Both datasets were then combined using the Unicycler assembly pipeline [33] with a finishing step of Pilon v1.22. The assemblies were visually inspected using the assembly graph tool Bandage v0.8.1 [34]. Annotation of the megaplasmid was performed with Prokka v1.13 using default parameters [35]. To improve annotation, we downloaded additional files of trusted proteins from NCBI RefSeq plasmids (ftp://ftp.ncbi.nih.gov/refseq/release/plasmid/), the NCBI Bacterial Antimicrobial Resistance Reference Gene Database (ftp://ftp.ncbi.nlm.nih.gov/pathogen/Antimicrobial_resistance/) and the Antibacterial Biocide- and Metal-Resistance Genes database (Bac-Met, http://bacmet.biomedicine.gu.se/index.html). EggNOG mapper v4.5.1 and NCBI’s Conserved Domain Database CDSEARCH/cdd v3.16 were used for functional annotation and conserved domain search of protein sequences, respectively [36–38]. Inference of orthologous groups (OGs) was achieved with OrthoFinder v2.2.6 [39]. The coding sequence (CDS) annotations of the megaplasmid were visualized with Circos v0.69-6 [40]. We used ISfinder [41] to look for insertion sequences (IS). Antimicrobial resistance genes and associated mobile elements were annotated using Galileo™ AMR (https://galileoamr.arcbio.com/mara/ (Arc Bio, Cambridge, MA) [42]. Plasmid copy number was estimated based on coverage of the Illumina dataset. GenSkew (http://genskew.csb.univie.ac.at/) was used to compute and plot nucleotide skew data to predict the origin of replication.

### Accession Number

The sequence of plasmid pJBCL41 was deposited in GenBank with accession number MK496050.

## Results

### Antimicrobial Susceptibility and Taxonomy testing

Clinical isolate FFUP_PS_41 has a multidrug resistance (MDR) phenotype, showing resistance to imipenem, meropenem, ceftazidime, cefepime, aztreonam, piperacilin+tazobactam, gentamicin, tobramycin, amikacin, ciprofloxacin but remains susceptible to colistin (MIC=1 mg/L). FFUP_PS_41 was initially identified as *P. putida* by VITEK-2. However, it displays an ANI value below the cut-off for species identification (95%) [28] when compared with the complete genome of type strains belonging to the *Pseudomonas* genus, suggesting that it represents a new species related to the *P. putida* phylogenetic group.

### Comparative Megaplasmidomics Between pJBCL41 and Related Pseudomonas Plasmids

Using a hybrid assembly approach, we were able to fully resolve a mosaic megaplasmid (named pJBCL41) carried by *Pseudomonas* sp. FFUP_PS_41 (**Figure S1**). pJBCL41 is 498,516 bp and a total of 608 predicted CDS were annotated (**Figure 1**). It has an average GC content of 56.0%, which is lower than that observed for the chromosome (62.6%) and the mean content for strains identified as *P. putida* (62.0%, according to information retrieved on the 08/03/2019 on https://www.ezbiocloud.net/taxon?tn=Pseudomonas%20putida).

**Figure 1.**
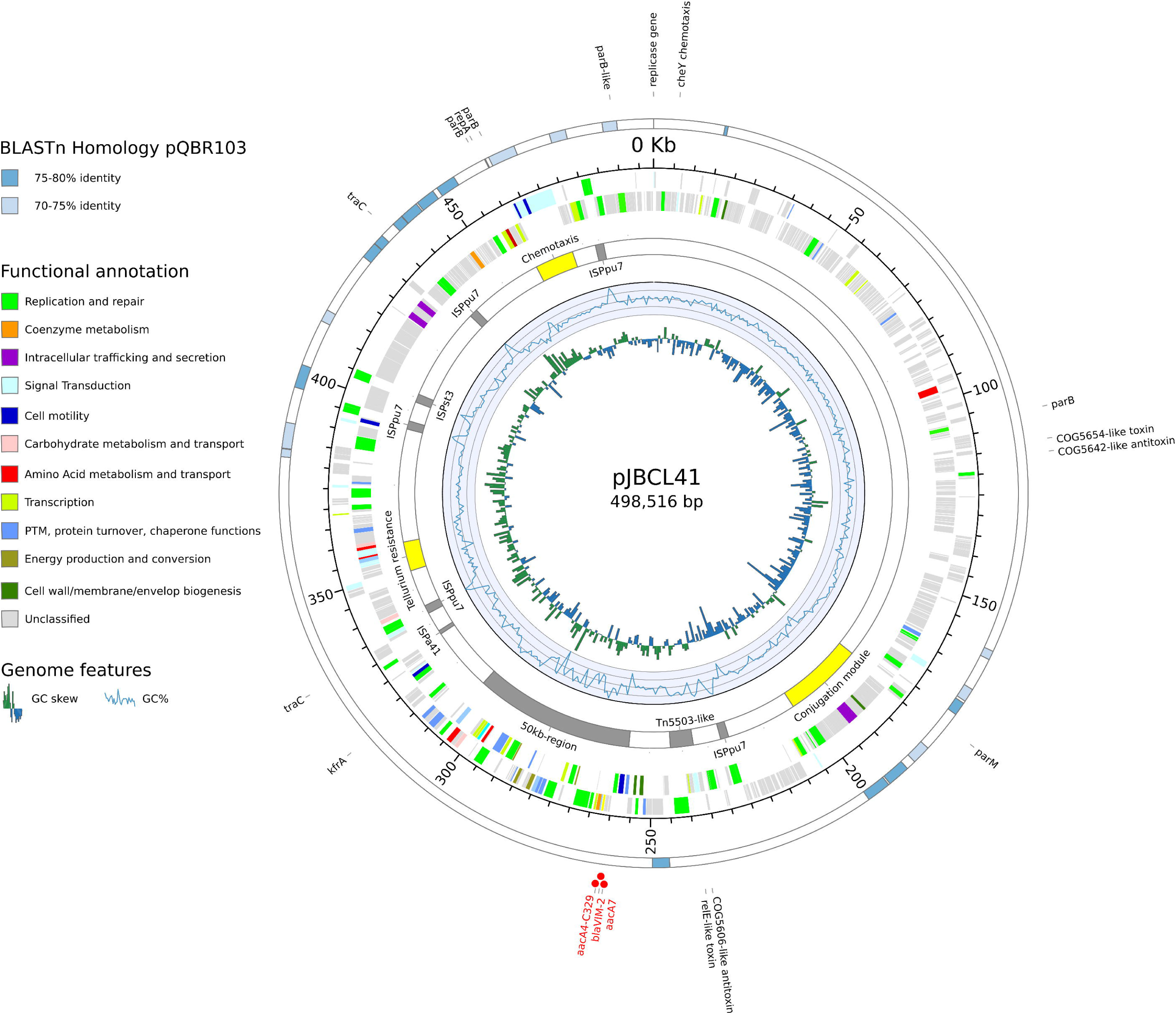
Circular representation of genomic features of pJBCL41. The innermost circle is a histogram of the GC skew, the next a graph of GC content. The next circle displays selected regions of interest (yellow) and IS and transposons or related elements (grey). Red dots highlight genes encoding for antibiotic resistance. The next two circles represent the coding regions on the negative and positive strands colored by their functional annotation (when available). The outermost circle displays regions with high levels of identity to pQBR103 (GenBank accession no. NC_009444.1).

NCBI’s CDD calls 42.1% (256) of the predicted CDS for pJBCL41 (**Table S1**), indicating that most genes encode proteins of unknown function. The backbone of this megaplasmid harbours genes predicted to be responsible for plasmid replication, heavy metal resistance and carries two predicted type-II toxin-antitoxin (TA) systems and genes encoding for partition systems (**Figure 1**) [43]. Several genes encoding transport and metabolic processes, as well as transposable elements and CDS associated with transcription, regulatory, chemotaxis signal transduction and mobility functions could be identified. These traits are frequently overrepresented on large plasmids (**Figure 2**) [6, 44]. Also, pJBCL41 harbours several genes coding for the synthesis of DNA precursors, which may promote replication and transcription processes to alleviate the burden that this acquired element may impose on the host cell.

**Figure 2.**
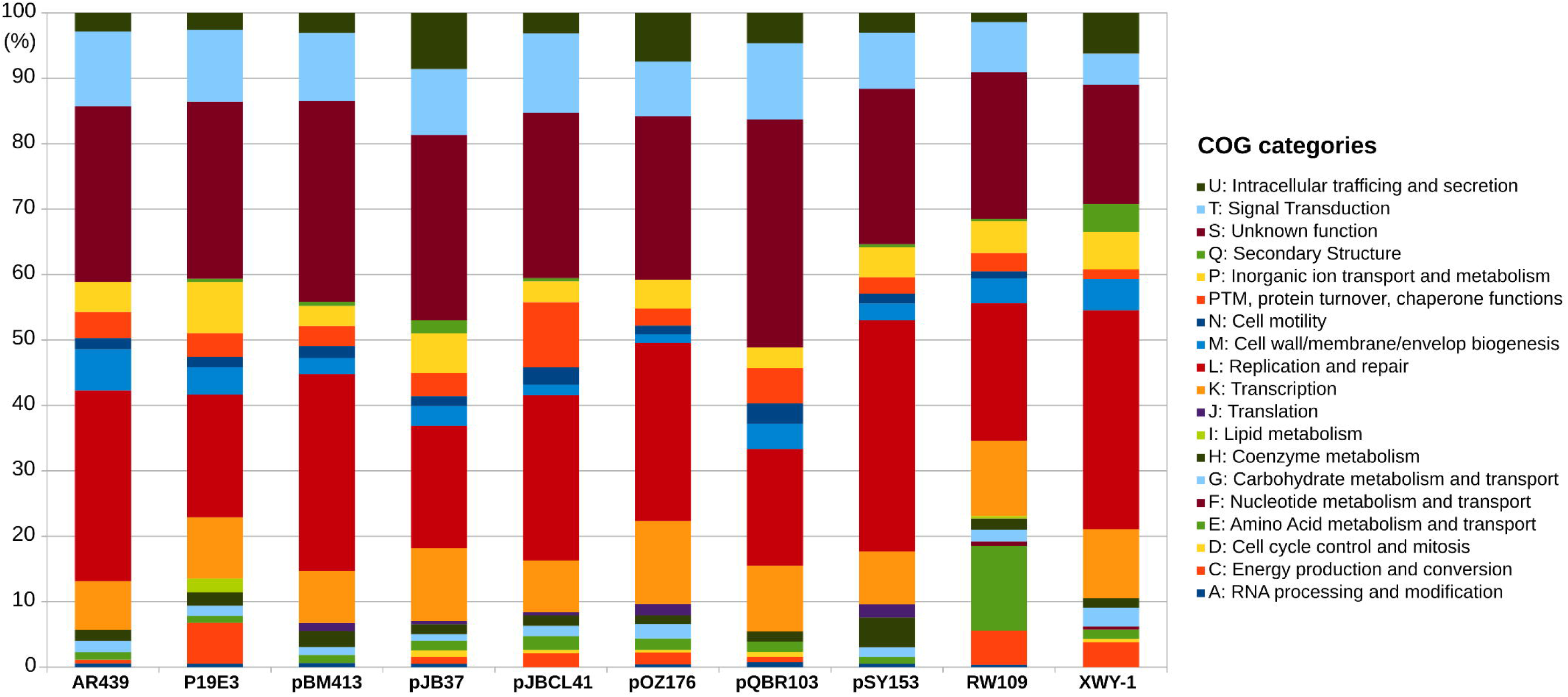
Functional characterization of pJBCL41 and related megaplasmids. COG stands for Cluster of Orthologous Groups.

pJBCL41 has low nucleotide sequence identity with *Pseudomonas* megaplasmids deposited in public databases (**Table 1** and **Figure S2**). OrthoFinder assigned 59.4% of proteins encoded by pJBCL41 and the most closely-related plasmid, pQBR103 from *Pseudomonas fluorescens* [45] into 335 OGs (**Table S2**). pQBR103 was found in *Pseudomonas* populations colonizing the leaf and root surfaces of sugar beet plants growing at Wytham, United Kingdom and carried no antimicrobial resistance genes [45]. Curiously, a blastp analysis between the proteins encoded by these megaplasmids revealed that the average amino acid sequence identity is 72.8% among sequences producing significant alignments.

**Table 1.**
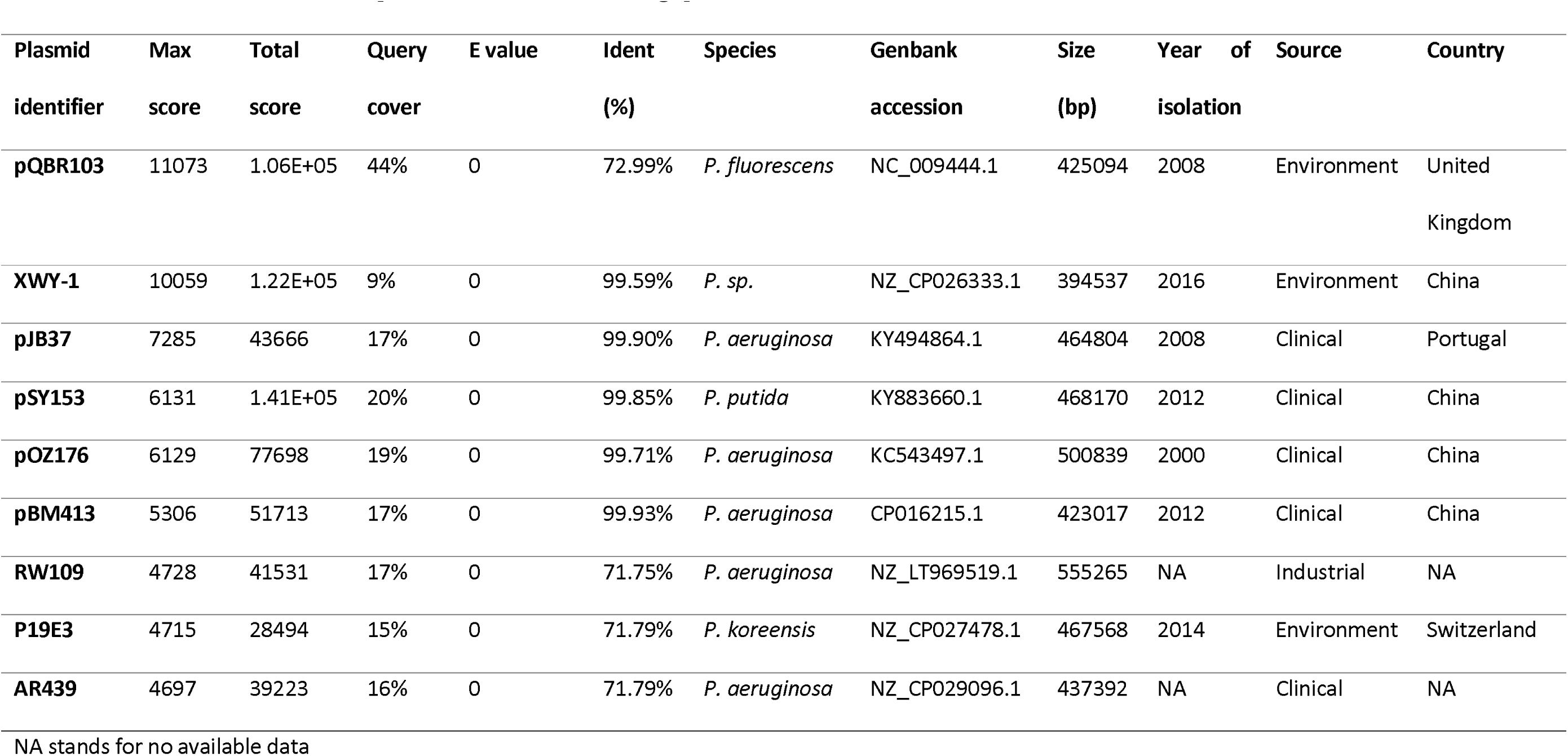
Blastn results between pJBCL41 and related megaplasmids.

Large plasmids identified among the *Pseudomonas* genus usually belong to the IncP-2 incompatibility group [10, 21, 24]. However, the IncP-2-type stability/replication/conjugal transfer system is absent from pJBCL41 as previously observed for other megaplasmids carried by different *Pseudomonas* species [46, 47]. Two replication proteins could be identified here. One replicase gene is located at 458,679 bp on the plasmid and is close to predicted the origin of replication (**Figure S3**). pJBCL41 is estimated to be present as a single copy, from read coverage vs. the chromosome. Like many megaplasmids, pJBCL41 appears to possess a full set of genes for self-transmission [2, 3]. We identified a cluster of genes encoding an F-type T4SS, encompassing i) a gene encoding a TraD homolog, an AAA+ ATPase of the pfamVirD4 type, known as the T4CP and which is a key protein in conjugation; ii) a gene encoding a TraI relaxase homolog, which together with accessory proteins is responsible for cleaving the plasmid in a site-specific manner to initiate DNA transfer and iii) a set of genes (*traEFGKNV* homologues) coding for the mating pair formation system responsible for pilus assembly and retraction (**Figure 1**) [2, 3, 48].

### pJBCL41 Carries a Complex 50 kb Multidrug Resistance Region

The plasmid pJBCL41 carries genes typically found on IncP-2 encoding resistance to tellurite, which could allow co-selection and enrichment of bacteria with MDR plasmids [49]. It also harbours a class 1 integron with the |*aacA7*|*bla*_VIM-2_|*aacA4*| cassette array (named In103 by INTEGRALL [50]) (**Figure 3**): *aacA7* confers resistance to aminoglycosides and *bla*_VIM-2_ encodes resistance to β-lactams (including carbapenems). The *aacA4* gene cassette has a C residue at nucleotide position 329, encoding a serine associated with gentamicin resistance [51]. The same cassette array has been observed previously among isolates from Portuguese hospitals [22]. The integron is of the In4 type, with a complete 5’-CS bounded by the 25 bp inverted repeat IRi, 2,239 bp of the 3’-CS and IS*6100* flanked by two fragments of the IRt end of Tn*402* [9, 52]. As the region between IRi and IRt lacks *tni* transposition genes, this constitutes a Tn*402*-like transposon that would be defective in self-transposition.

**Figure 3.**
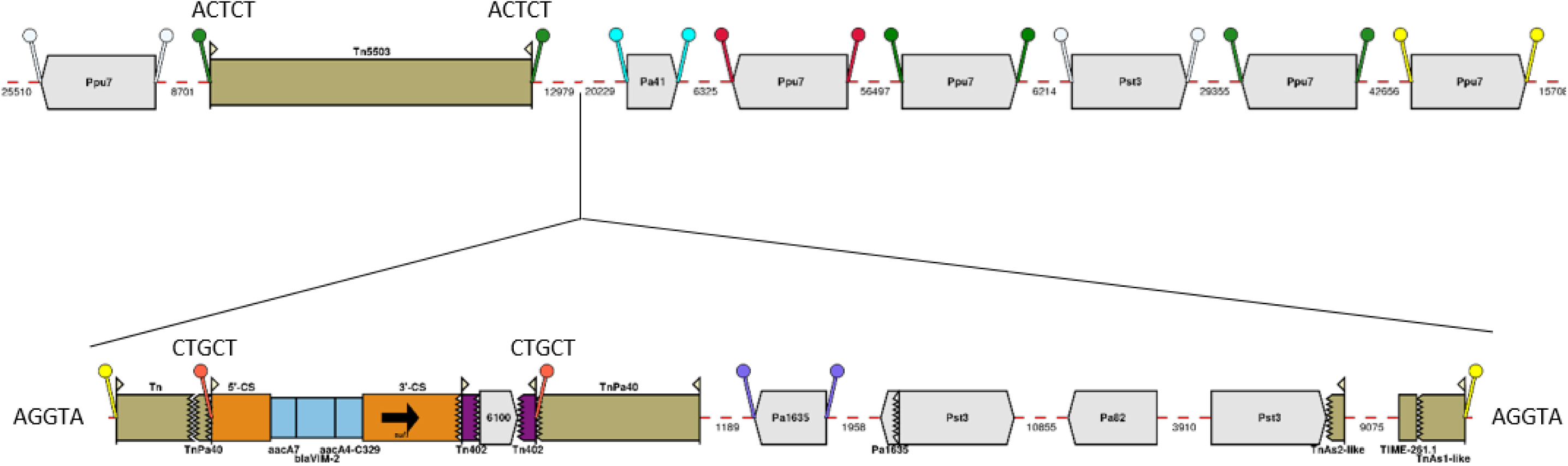
Map of resistance genes and mobile genetic elements inserted in the backbone of pJBLC41. Gene cassettes are shown as blue boxes labelled with the cassette name and are oriented in the 5’-CS to 3’-CS direction. IS are shown as block arrows labelled with the IS name/number, with the pointed end corresponding to IR_R_. TIME-261.1 and fragments of Tn*3*-family transpospons are shown as beige boxes with 38 bp IR represented by flags. The fragment annotated as “TnAs1-like” is ∼97% identical to a region in common between Tn*1721* (GenBank accession no. X61367.1) and Tn*As1* in ISfinder. The fragment annotated as “TnAs2-like” is ∼94% identical to Tn*As2* in ISfinder. The integron is inserted in a proposed hybrid transposon, apparently created by *res*-mediated recombination between a *tnp* region matching Tn*Pa40* and another tranpsoson, labelled “Tn”, that is ∼86% identical to Tn*As1* over the ∼300 bp at the IR_L_ end only. Direct repeats are shown as a pair of ‘lollipops’ of the same colour flanking an IS or a pair of IRs (but note that the same colour may be used to indictate more than one pair of DR), with sequences indicated for DR of transposons. Mobile elements are shown to scale and numbers below dashed red lines indicate the lengths of intervening regions in bp. This figure was constructed from diagrams generated using Galileo™ AMR.

This defective Tn*402*-like transposon is flanked by 5-bp direct repeats (5’-CTGCT-3’) (**Figure 3**), suggesting integration by transposition close to the predicted resolution (*res*) site of a Tn*3*-family transposon. About 300 bp at the IR_L_ end of the transposon are related (∼86% identical) to Tn*As1* (ISfinder), followed by a region containing a gene which may encode a methyl-accepting chemotaxis protein. From the predicted recombination crossover point in the *res* site the sequence matches Tn*Pa40* (ISfinder). This “hybrid” transposon is not flanked by characteristic 5 bp DR but the 5 bp adjacent to IR_L_ (5’-AGGTA-3’) are repeated 50,273 bp away, immediately adjacent to the 38 bp repeat of a 1,100 bp transposon fragment ∼97% identical to part of both Tn*1721* (GenBank accession no. X61367.1, [53]) and Tn*As1* (**Figure 3**). This transposon is truncated by 261 bp region that apparently corresponds to a Tn*3*-Derived Inverted-Repeat Miniature Element (designated TIME-262.1 here). TIMEs are non-autonomous mobile elements commonly found in *Pseudomonas* spp. [54].

Most of the region between these transposon elements consists of a 16,782 bp segment flanked by directly oriented copies of IS*Pst3* (IS*21* family). This region, except for insertion of IS*Pa82* (IS*66* family) and an adjacent deletion in pJBCL41, matches several *Pseudomonas* chromosomes (e.g. *P. aeruginosa* PA7 in **Figure S4**) and different parts of it are found in plasmids in *Pseudomonas, Acinetobacter* and *Enterobacteriaceae*, sometimes also flanked by IS. The sequence between Tn*Pa40* and the left-hand IS*Pst3* in pJBCL41 is a duplication of part of the 16,782 bp region, with IS*Pa1635* (IS*4* family) inserted, flanked by characteristic 8 bp DR, instead of IS*Pa82* and ends with a partial IS*Pa1635*. The right-hand IS*Pst3* truncates a transposon related to Tn*As2* [55], which is separated from TIME-261.1 by a 9,075 bp region that also matches *Pseudomonas* chromosomes and includes a putative aminoglycoside phosphotransferase gene.

Blast searches with the complete 50 kb region identified a 59 kb region in the chromosome of *P. aeruginosa* AR_0440 (GenBank accession no. CP029148.1) that has similar ends, but lacks an integron, with an additional Tn*5393* insertion and a different region in place of the IS*Pst3*-bounded segment (**Figure S4**). This 59 kb region is flanked by 5 bp DR (5’-AATGA-3’) and an uninterrupted version of the flanking sequence matches other *Pseudomonas* chromosomes.

A Tn*5503*-like transposon encoding a type-II TA system and two metal dependent phosphohydrolases is also inserted in pJBCL41 [56] and is flanked by 5-bp DR (5’-ACTCT-3’), indicating that this element transposed independently of the 50-kb region (**Figure 3**). It has only 10 nucleotide differences from the original Tn*5503* on plasmid Rms149, the archetype of *Pseudomonas* plasmid incompatibility group IncP-6 [56], and additional copies of short repeats in a GC-rich region within a gene encoding an ATP-utilizing enzyme. An additional IS*Pst3*, five IS*Ppu7* (IS*21* family) and one IS*Pa41* (IS5 family) all flanked by DR of characteristic length, are also inserted in the pJBCL41 backbone (**Figures 1 and 3**).

## Discussion

In this study, we took advantage of a hybrid assembly approach to fully resolve and characterize a carbapenemase-encoding megaplasmid and to compare it with related *Pseudomonas* megaplasmids. The lower GC content of pJBCL41 compared with the FFUP_PS_41 chromosome and strains identified as *P. putida* may be related to a more relaxed selection acting on these secondary replicons, as the maintenance of GC-rich genomes is energetically more demanding [57, 58]. Ongoing studies will help to characterize the biology and genomic signatures related to this new putative *Pseudomonas* species (Botelho *et al*, unpublished data).

Since secondary replicons are under strong pressure to undergo genomic reshuffling [57], the observed low nucleotide sequence identity between pJBCL41 megaplasmids and large *Pseudomonas* plasmids deposited in public databases might be expected. Even though pJBCL41 and pQBR103 plasmids are similar in size and functionalities, there is a high level of divergence between genes encoding related proteins. Indeed, it is rare to identify megaplasmids with a similar nucleotide sequence in strains belonging to different species within the same genus [6, 47]. These results suggest that pJBCL41 and pQBR103 may share a common ancestor, but independent evolutionary trajectories have led to significant diversification among related genes. The presence of different replicons suggests that pJBCL41 may have resulted from co-integration of distinct plasmid modules. The replication module defines plasmid copy number and plasmid survival in several hosts. Low copy-number plasmids are more frequently lost, due to random assortment at cell division [2, 3] and extra stability modules, such as TA and partition systems, may be required to ensure that large plasmids such as pJBCL41 are maintained [43, 59].

The DR flanking the 50-kb region in pJBCL41 and the related 59-kb region in the *P. aeruginosa* AR_0440 chromosome could reflect insertion of each region by transposition, possibly mediated by the intact transposase and resolvase of Tn*Pa40*.

However, the size, complexity and differences the internal parts of these related regions may be more consistent with initial insertion of a simple transposon followed by further insertions, deletions and rearrangements. A similar situation is seen in plasmid pCTX-M360, which carries a complete Tn*2* flanked by the 5 bp DR, and the highly-related pCTX-M3, in which the ends of Tn*2* are present in the same position but the central part of the transposon has undergone extensive rearrangements [60]. The identification of all or part of the 16,782 bp segment found within the 50 kb region in pJBCL41 in other locations also suggests that some of the genes it carries may encode advantageous functions, but this needs further analysis. Identification of other sequences related to parts of these 50-kb and 59-kb region segments may also shed light on how they have arisen and evolved.

To sum up, we show that a hybrid Nanopore/Illumina approach is useful for producing contiguous assemblies and allowed full resolution of a carbapenemase-encoding *Pseudomonas* megaplasmid. The presence of this large plasmid may provide a selective advantage to the host cell. However, given their size and gene content, acquisition of these secondary replicons may pose a significant cost [61–63]. The high level of gene variation when compared to publicly available megaplasmids suggests that these secondary replicons frequently undergo gene loss and gain though HGT. The reduced purifying selection and the high prevalence of transposable elements frequently observed on megaplasmids may help to explain why these elements readily acquire foreign DNA [6, 57, 64]. In fact, mosaic plasmids such as pJBCL41 and the majority of megaplasmids have a high proportion of mobile genetic elements [8]. The identification of novel megaplasmids may shed light on the evolutionary effects of gene transfer and the selective forces driving antibiotic resistance.

## Acknowledgments

This work was supported by the Applied Molecular Biosciences Unit- UCIBIO which is financed by national funds from FCT/MCTES (UID/Multi/04378/2019). JB and FG were supported by grants from Fundação para a Ciência e a Tecnologia (SFRH/BD/104095/2014 and SFRH/BPD/95556/2013, respectively). CL is supported by an SB PhD fellowship from FWO Vlaanderen (1S64718N).

