## Supplemental Figure 1 for "Megaplasmids on the Rise: Combining Sequencing Approaches to Fully Resolve a Carbapenemase-Encoding Plasmid in a Proposed Novel *Pseudomonas* Species"

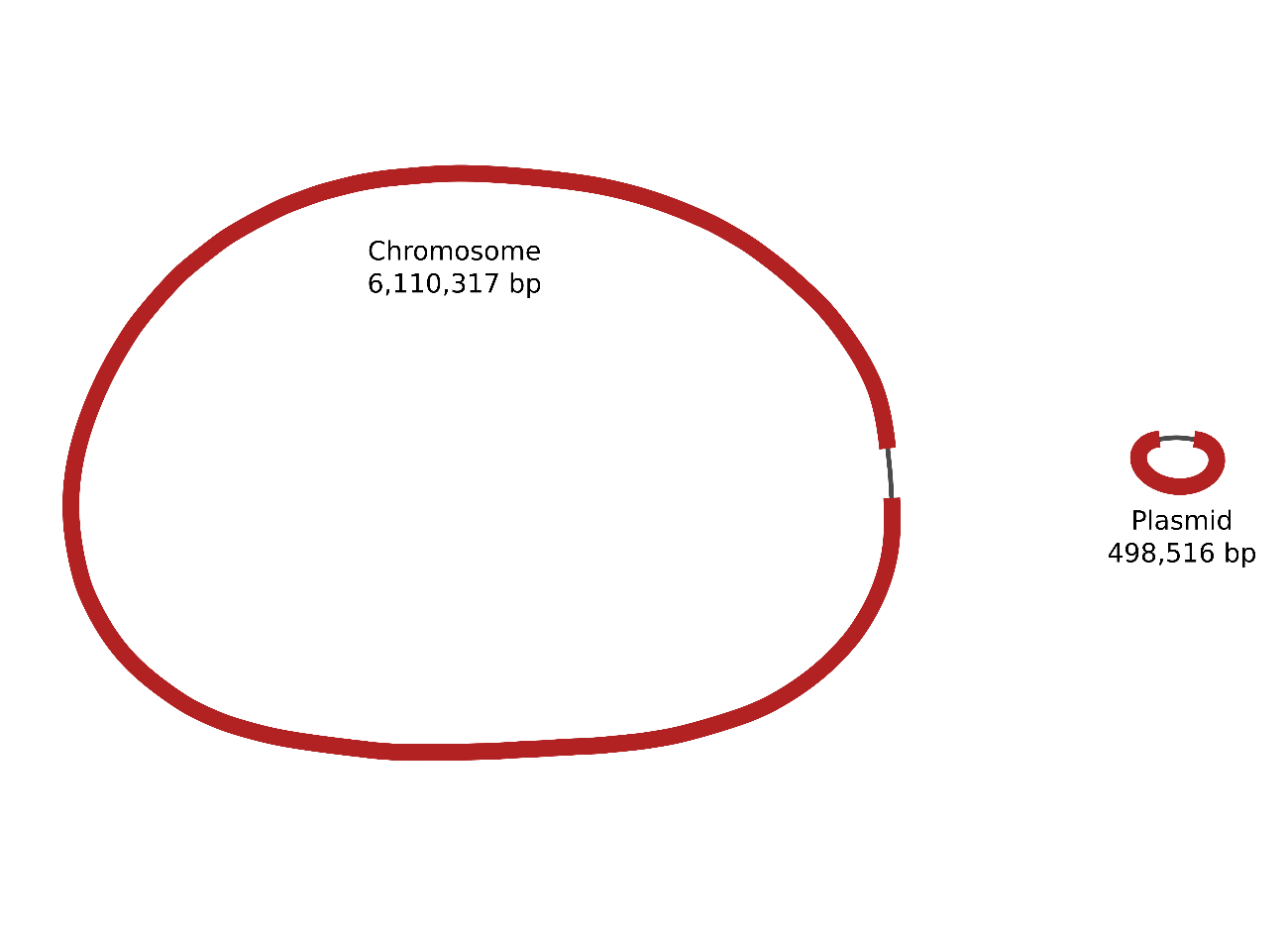

**Figure S1** – Visual representation of *de novo* assembly graph of strain FFUP_PS_41 containing the pJBCL41 megaplasmid using Bandage.
