## Supplemental Figure 2 for "Megaplasmids on the Rise: Combining Sequencing Approaches to Fully Resolve a Carbapenemase-Encoding Plasmid in a Proposed Novel *Pseudomonas* Species"

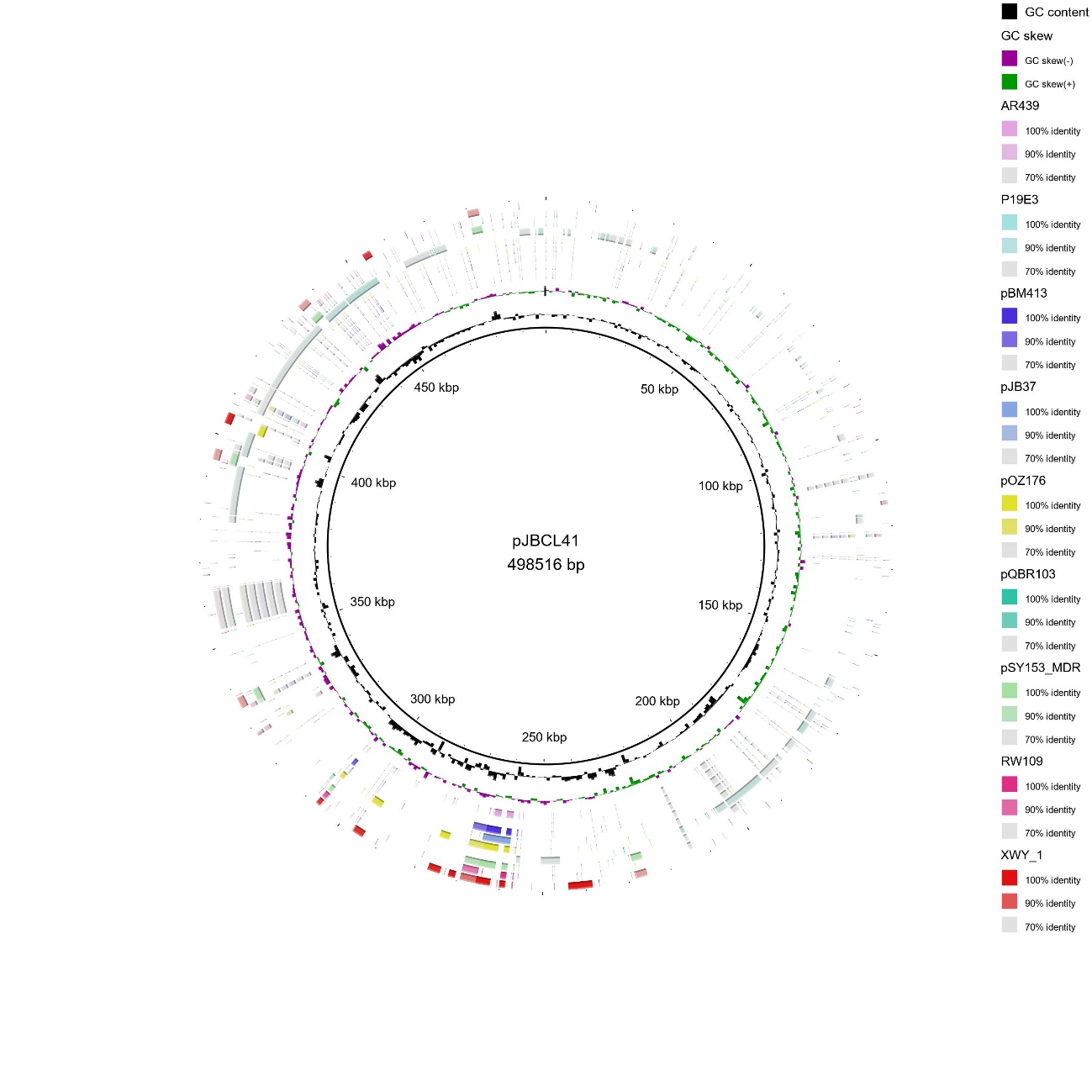

**Figure S2**. Comparison between the complete nucleotide sequence of pJBCL41 and related *Pseudomonas* megaplasmids.
