## Supplementary figures and images for "Megaplasmids on the Rise: Combining Sequencing Approaches to Fully Resolve a Carbapenemase-Encoding Plasmid in a Proposed Novel *Pseudomonas* Species"

### Supplemental Figure 3

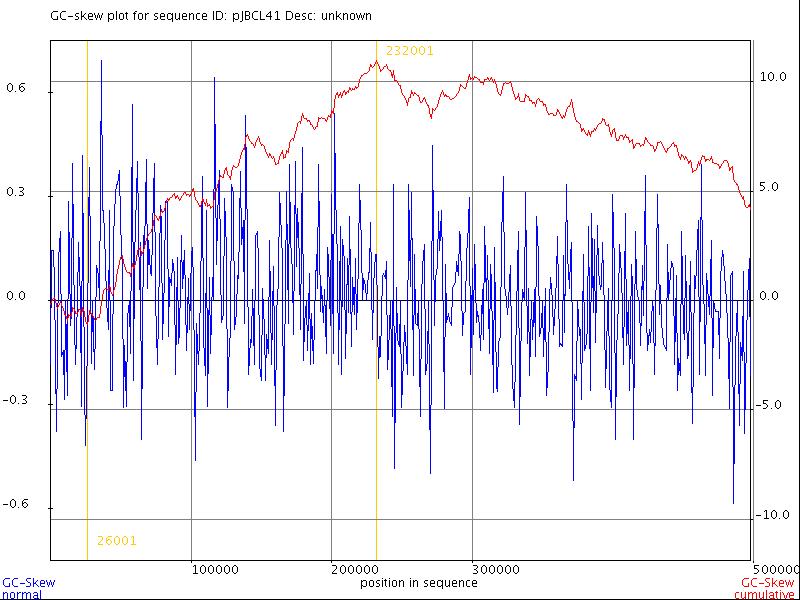

**Figure S3**. Origin (26,001-bp) and terminus (232,001-bp) of DNA replication using GC skew and cumulative GC skew plots.
