## Supplemental Figure 4 for "Megaplasmids on the Rise: Combining Sequencing Approaches to Fully Resolve a Carbapenemase-Encoding Plasmid in a Proposed Novel *Pseudomonas* Species"

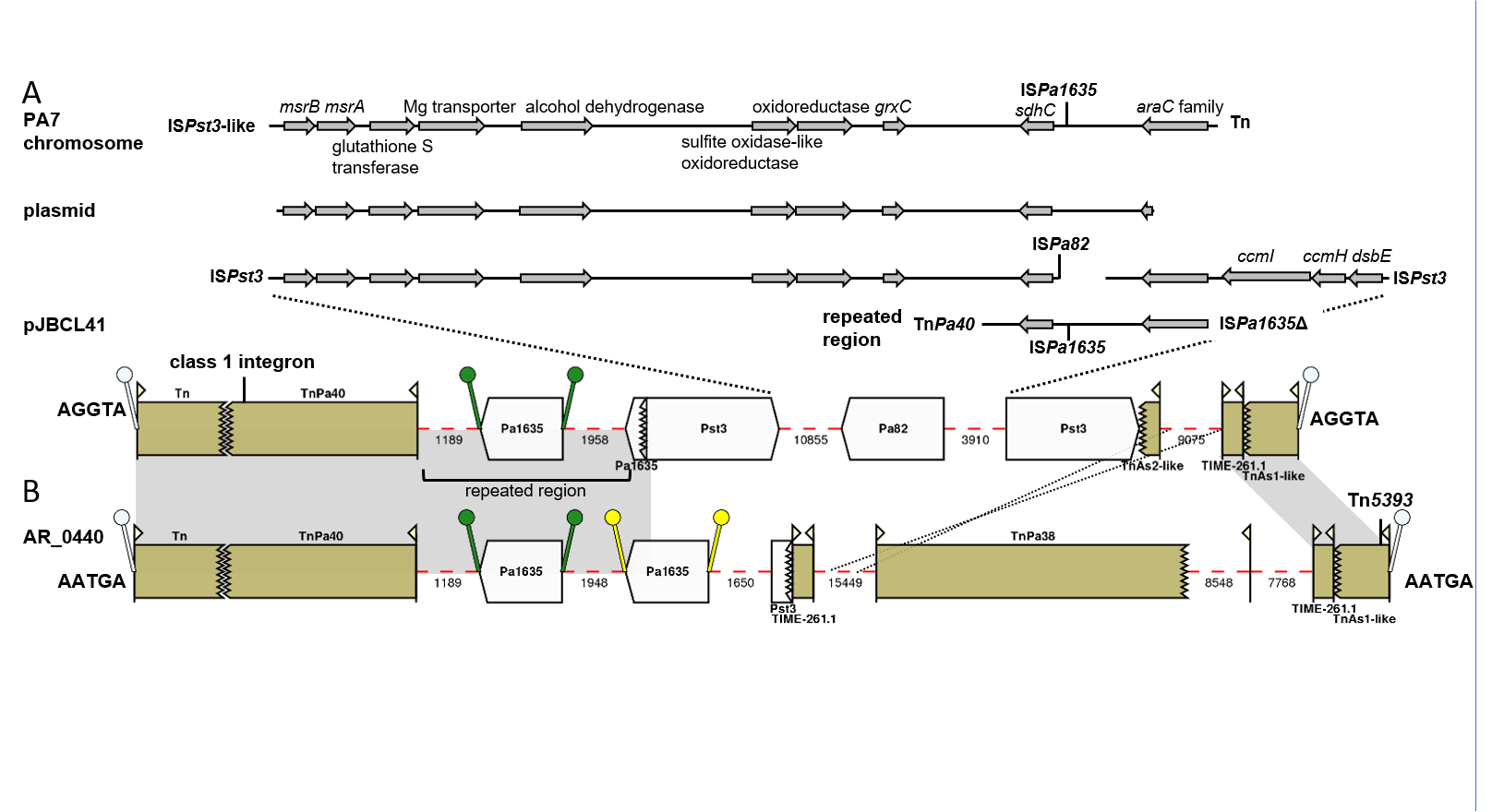

**Figure S4. A.** Comparison of the IS*Pst3*-flanked region in pJBCL41 with part of the *P. aeruginosa* PA7 chromosome (GenBank accession no. CP000744.1) and a representative plasmid (pCAV2018-177, GenBank accession no. CP029430.1). Inserted IS are shown above (all IS*Pa1635* copies have flanking DR, IS*Pa82* does not) and the gap adjacent to IS*Pa82* in pJBCL41 indicates a deletion. The names of IS and Tn defining the ends of the matching regions are shown adjacent to these regions. Predicted gene names/functions of selected orfs are shown (additional hypothetical proteins are not marked). The region between Tn*Pa40* and IS*Pa1635*Δ in pJBCL41 that forms a partial repeat is also shown, **B.** Comparison of the 50 kb insertion in pJBCL41 with the 59 kb insertion in the chromosome of *P. aeruginosa* AR_0440 (GenBank accession no. CP029148.1). Diagrams were generated using Galileo^TM^ AMR, with mobile elements shown to scale and numbers below dashed red lines indicating the lengths of intervening regions in bp. Matching regions are indicated by grey shading. The 9,075 bp region between the Tn*As2*-like transposon and TIME-261.1 in pJBCL41 matches positions 9836-762 of the 15,449 bp region between TIME-261.1 and Tn*Pa38* in AR_0440, shown by dotted lines. DR are shown by white ‘lollipops’ and sequences are indicated. The position of the class 1 integron in pJBCL41 and Tn*5393* in AR_0440, both flanked by 5 bp DR, are indicated by labelled vertical lines.
