## Supplemental Table 1 for "Megaplasmids on the Rise: Combining Sequencing Approaches to Fully Resolve a Carbapenemase-Encoding Plasmid in a Proposed Novel *Pseudomonas* Species"

**Table S1**. NCBI’s CDD annotation of protein sequences from the pJBCL41 megaplasmid.

| **Locus tag** | **Hit type** | **PSSM-ID** | **From** | **To** | **E-Value** | **Bitscore** | **Accession** | **Short name** | **Incomplete** | **Superfamily** |
| --- | --- | --- | --- | --- | --- | --- | --- | --- | --- | --- |
| pJBCL41_00001 | specific | 234973 | 1 | 58 | 2.61049e-29 | 97.6714 | PRK01712 | PRK01712 | NA | cl00670 |
| pJBCL41_00005 | superfamily | 295372 | 16 | 77 | 0.0096178 | 32.6895 | cl02570 | RhoGAP superfamily | NC | NA |
| pJBCL41_00006 | specific | 234726 | 1 | 304 | 2.71791e-131 | 374.947 | PRK00321 | rdgC | NA | cl01122 |
| pJBCL41_00007 | superfamily | 328931 | 13 | 55 | 1.3546e-07 | 45.2864 | cl22854 | HTH_XRE superfamily | C | NA |
| pJBCL41_00007 | specific | 316117 | 74 | 107 | 0.000413435 | 35.9761 | pfam13560 | HTH_31 | C | cl22854 |
| pJBCL41_00013 | specific | 238088 | 9 | 123 | 1.97283e-33 | 112.669 | cd00156 | REC | NA | cl19078 |
| pJBCL41_00015 | specific | 223861 | 57 | 179 | 1.50269e-06 | 48.5019 | COG0790 | TPR | NC | cl27881 |
| pJBCL41_00016 | superfamily | 329030 | 10 | 198 | 8.35574e-17 | 78.4374 | cl23717 | crotonase-like superfamily | NA | NA |
| pJBCL41_00021 | specific | 223394 | 8 | 153 | 3.78905e-17 | 77.6759 | COG0317 | SpoT | C | cl28493 |
| pJBCL41_00023 | superfamily | 321445 | 39 | 127 | 0.00534522 | 35.3357 | cl01317 | Cmr5_III-B superfamily | C | NA |
| pJBCL41_00026 | superfamily | 330238 | 6 | 438 | 3.66699e-87 | 274.661 | cl25417 | CDC9 superfamily | NA | NA |
| pJBCL41_00028 | specific | 225650 | 82 | 194 | 1.05368e-24 | 95.9001 | COG3108 | YcbK | N | cl01194 |
| pJBCL41_00029 | specific | 143586 | 114 | 217 | 5.06998e-40 | 132.677 | cd07185 | OmpA_C-like | NA | cl28145 |
| pJBCL41_00030 | superfamily | 329113 | 59 | 138 | 2.52256e-06 | 45.6345 | cl23836 | DUF262 superfamily | C | NA |
| pJBCL41_00042 | superfamily | 329030 | 78 | 209 | 2.03213e-16 | 77.667 | cl23717 | crotonase-like superfamily | N | NA |
| pJBCL41_00046 | specific | 238846 | 23 | 212 | 1.34557e-24 | 98.631 | cd01713 | PAPS_reductase | NA | cl00292 |
| pJBCL41_00052 | specific | 307387 | 15 | 82 | 1.53442e-22 | 81.8251 | pfam01206 | TusA | NA | cl00436 |
| pJBCL41_00053 | specific | 311218 | 2 | 72 | 2.74317e-20 | 76.4242 | pfam07130 | YebG | NA | cl01217 |
| pJBCL41_00054 | specific | 223363 | 71 | 254 | 1.29375e-05 | 45.8705 | COG0286 | HsdM | NC | cl28090 |
| pJBCL41_00059 | superfamily | 320736 | 37 | 122 | 0.000564712 | 39.3582 | cl00057 | vWFA superfamily | N | NA |
| pJBCL41_00060 | superfamily | 330529 | 55 | 157 | 0.00156293 | 37.7391 | cl25708 | Peptidase_M16 superfamily | C | NA |
| pJBCL41_00060 | superfamily | 332840 | 19 | 52 | 0.00568678 | 33.348 | cl28019 | Arc superfamily | C | NA |
| pJBCL41_00063 | superfamily | 318666 | 11 | 96 | 0.0053245 | 33.9815 | cl24921 | KORA superfamily | NA | NA |
| pJBCL41_00069 | specific | 223627 | 89 | 521 | 1.17479e-17 | 86.7124 | COG0553 | HepA | N | cl26465 |
| pJBCL41_00072 | superfamily | 333066 | 2 | 66 | 1.00894e-20 | 86.3517 | cl28246 | DnaJ superfamily | C | NA |
| pJBCL41_00081 | superfamily | 329065 | 65 | 143 | 0.00116487 | 37.6913 | cl23771 | Big_3_5 superfamily | NA | NA |
| pJBCL41_00089 | specific | 235838 | 9 | 291 | 4.23819e-153 | 428.825 | PRK06596 | PRK06596 | NA | cl27146 |
| pJBCL41_00094 | superfamily | 332840 | 11 | 55 | 5.02763e-13 | 58.3859 | cl28019 | Arc superfamily | NA | NA |
| pJBCL41_00112 | specific | 312300 | 18 | 159 | 5.2187e-52 | 163.111 | pfam08719 | DUF1768 | NA | cl21532 |
| pJBCL41_00114 | superfamily | 330277 | 1 | 194 | 6.09235e-22 | 88.2281 | cl25456 | PRK15113 superfamily | NA | NA |
| pJBCL41_00124 | specific | 306954 | 41 | 145 | 1.0179e-12 | 62.1422 | pfam00583 | Acetyltransf_1 | NA | cl17182 |
| pJBCL41_00128 | specific | 197273 | 15 | 115 | 3.52584e-13 | 65.442 | cd09176 | PLDc_unchar6 | NA | cl15239 |
| pJBCL41_00136 | specific | 308282 | 9 | 209 | 7.44671e-46 | 150.755 | pfam02586 | SRAP | NA | cl03646 |
| pJBCL41_00138 | specific | 308235 | 132 | 218 | 9.12701e-05 | 39.8637 | pfam02517 | Abi | NA | cl00558 |
| pJBCL41_00152 | superfamily | 320882 | 51 | 378 | 3.34454e-31 | 126.39 | cl00292 | AANH_like superfamily | C | NA |
| pJBCL41_00159 | superfamily | 329065 | 72 | 160 | 7.15356e-05 | 40.7729 | cl23771 | Big_3_5 superfamily | NA | NA |
| pJBCL41_00164 | superfamily | 320982 | 46 | 80 | 0.00451195 | 33.8002 | cl00456 | SLC5-6-like_sbd superfamily | NC | NA |
| pJBCL41_00165 | superfamily | 331543 | 152 | 206 | 0.000434074 | 40.4403 | cl26722 | Spo0J superfamily | N | NA |
| pJBCL41_00176 | specific | 143393 | 98 | 158 | 2.68626e-09 | 51.6487 | cd05403 | NT_KNTase_like | C | cl11966 |
| pJBCL41_00177 | superfamily | 332840 | 37 | 86 | 5.60129e-06 | 42.2076 | cl28019 | Arc superfamily | NA | NA |
| pJBCL41_00178 | specific | 238038 | 245 | 301 | 6.36943e-07 | 45.5404 | cd00085 | HNHc | NA | cl00083 |
| pJBCL41_00180 | superfamily | 321922 | 13 | 163 | 2.94304e-33 | 115.245 | cl02411 | RES superfamily | NA | NA |
| pJBCL41_00181 | superfamily | 327431 | 47 | 168 | 2.16967e-20 | 81.3484 | cl17718 | DUF2384 superfamily | NA | NA |
| pJBCL41_00186 | superfamily | 330239 | 1 | 273 | 2.30051e-99 | 309.143 | cl25418 | ligD superfamily | NC | NA |
| pJBCL41_00190 | superfamily | 327401 | 19 | 240 | 6.81649e-83 | 250.707 | cl17173 | AdoMet_MTases superfamily | NA | NA |
| pJBCL41_00190 | superfamily | 330753 | 262 | 318 | 0.00542513 | 38.1452 | cl25932 | Peptidase_S21 superfamily | N | NA |
| pJBCL41_00196 | superfamily | 317636 | 125 | 185 | 3.81131e-06 | 45.939 | cl25918 | CCDC14 superfamily | NC | NA |
| pJBCL41_00198 | superfamily | 325091 | 22 | 133 | 2.48597e-10 | 54.0111 | cl11584 | DUF2591 superfamily | NA | NA |
| pJBCL41_00203 | superfamily | 320874 | 45 | 119 | 0.00932019 | 34.5522 | cl00276 | Maf_Ham1 superfamily | C | NA |
| pJBCL41_00226 | specific | 315954 | 319 | 368 | 6.10765e-17 | 74.3291 | pfam13391 | HNH_2 | NA | cl00083 |
| pJBCL41_00226 | superfamily | 225973 | 255 | 367 | 3.79898e-05 | 45.1923 | cl27449 | COG3440 superfamily | N | NA |
| pJBCL41_00228 | specific | 313019 | 9 | 42 | 0.0035941 | 31.0571 | pfam09723 | Zn-ribbon_8 | NA | cl00993 |
| pJBCL41_00235 | superfamily | 113356 | 286 | 395 | 0.0053284 | 38.7458 | cl25575 | Reo_sigmaC superfamily | C | NA |
| pJBCL41_00237 | superfamily | 331068 | 161 | 223 | 0.000513471 | 42.0359 | cl26247 | DNA_pol3_delta2 superfamily | NC | NA |
| pJBCL41_00244 | specific | 312369 | 35 | 183 | 9.87742e-15 | 68.6732 | pfam08808 | RES | NA | cl02411 |
| pJBCL41_00254 | specific | 236978 | 4 | 333 | 6.10103e-150 | 424.628 | PRK11778 | PRK11778 | NA | cl26733 |
| pJBCL41_00261 | superfamily | 330857 | 228 | 518 | 1.21941e-57 | 197.545 | cl26036 | YesN superfamily | N | NA |
| pJBCL41_00262 | superfamily | 324356 | 46 | 117 | 0.00431426 | 39.0222 | cl09326 | MATE_like superfamily | NC | NA |
| pJBCL41_00263 | superfamily | 333130 | 85 | 211 | 0.00174808 | 38.7315 | cl28310 | Rho superfamily | C | NA |
| pJBCL41_00265 | superfamily | 331610 | 88 | 405 | 1.7537e-11 | 65.32 | cl26789 | Toprim_N superfamily | NA | NA |
| pJBCL41_00271 | specific | 212669 | 8 | 337 | 1.15779e-39 | 142.171 | cd10227 | ParM_like | NA | cl17037 |
| pJBCL41_00279 | superfamily | 320948 | 140 | 252 | 0.000860654 | 39.566 | cl00388 | Thioredoxin_like superfamily | N | NA |
| pJBCL41_00281 | specific | 238157 | 139 | 202 | 1.43331e-05 | 42.7791 | cd00254 | LT_GEWL | C | cl00222 |
| pJBCL41_00281 | superfamily | 320832 | 10 | 58 | 0.000487475 | 39.2948 | cl00222 | lysozyme_like superfamily | C | NA |
| pJBCL41_00283 | superfamily | 331285 | 106 | 364 | 0.000343872 | 43.0033 | cl26464 | Atrophin-1 superfamily | N | NA |
| pJBCL41_00284 | superfamily | 330976 | 3 | 54 | 0.00204554 | 35.9911 | cl26155 | BamD superfamily | C | NA |
| pJBCL41_00285 | superfamily | 331107 | 471 | 927 | 5.7553e-46 | 177.518 | cl26286 | VirB4 superfamily | N | NA |
| pJBCL41_00287 | superfamily | 327560 | 25 | 129 | 6.72976e-08 | 48.0041 | cl19474 | TraV superfamily | NA | NA |
| pJBCL41_00288 | superfamily | 330184 | 291 | 501 | 2.11549e-09 | 57.1312 | cl25363 | VirB10_like superfamily | NA | NA |
| pJBCL41_00288 | superfamily | 328214 | 65 | 125 | 0.000284156 | 42.5644 | cl20817 | GBP_C superfamily | N | NA |
| pJBCL41_00289 | superfamily | 323239 | 30 | 207 | 6.37045e-05 | 42.714 | cl05878 | TraK superfamily | NA | NA |
| pJBCL41_00290 | superfamily | 322925 | 19 | 155 | 0.0044485 | 36.4725 | cl05060 | TraE superfamily | C | NA |
| pJBCL41_00295 | superfamily | 328724 | 130 | 556 | 1.46582e-19 | 92.3389 | cl21455 | P-loop_NTPase superfamily | N | NA |
| pJBCL41_00297 | superfamily | 331083 | 23 | 725 | 1.36665e-24 | 109.488 | cl26262 | RecD superfamily | NA | NA |
| pJBCL41_00299 | specific | 259853 | 2 | 87 | 2.97027e-40 | 127.507 | cd13831 | HU | NA | cl00257 |
| pJBCL41_00302 | specific | 224648 | 17 | 134 | 2.0796e-22 | 85.1472 | COG1734 | DksA | NA | cl00755 |
| pJBCL41_00305 | specific | 176454 | 9 | 352 | 3.33164e-174 | 490.137 | cd01700 | PolY_Pol_V_umuC | NA | cl25410 |
| pJBCL41_00305 | specific | 316001 | 375 | 422 | 1.68342e-15 | 69.8078 | pfam13438 | DUF4113 | NA | cl16275 |
| pJBCL41_00306 | specific | 119397 | 55 | 134 | 4.27904e-22 | 83.3772 | cd06529 | S24_LexA-like | NA | cl10465 |
| pJBCL41_00310 | specific | 239905 | 13 | 74 | 1.58059e-13 | 63.7518 | cd04458 | CSP_CDS | NA | cl09927 |
| pJBCL41_00312 | specific | 178993 | 1 | 332 | 0 | 538.681 | PRK00378 | PRK00378 | NA | cl15473 |
| pJBCL41_00314 | superfamily | 325091 | 6 | 113 | 3.09937e-11 | 55.9371 | cl11584 | DUF2591 superfamily | NA | NA |
| pJBCL41_00319 | specific | 225718 | 212 | 364 | 1.38853e-14 | 74.7932 | COG3177 | COG3177 | NC | cl26082 |
| pJBCL41_00325 | superfamily | 316032 | 40 | 89 | 0.00133807 | 36.9334 | cl24259 | Transglut_core3 superfamily | N | NA |
| pJBCL41_00328 | superfamily | 328724 | 54 | 229 | 3.06411e-74 | 223.345 | cl21455 | P-loop_NTPase superfamily | NA | NA |
| pJBCL41_00328 | specific | 312099 | 24 | 51 | 2.68876e-10 | 53.9011 | pfam08483 | IstB_IS21_ATP | NA | cl26739 |
| pJBCL41_00329 | specific | 226950 | 34 | 289 | 5.30137e-39 | 142.981 | COG4584 | COG4584 | NA | cl27787 |
| pJBCL41_00329 | specific | 226950 | 229 | 499 | 5.99373e-15 | 74.8007 | COG4584 | COG4584 | NA | cl27787 |
| pJBCL41_00334 | specific | 281099 | 182 | 465 | 5.91671e-119 | 350.779 | pfam03050 | DDE_Tnp_IS66 | NA | cl24150 |
| pJBCL41_00334 | specific | 225970 | 100 | 258 | 8.51109e-35 | 127.563 | COG3436 | COG3436 | NA | cl26266 |
| pJBCL41_00334 | specific | 316346 | 471 | 507 | 1.48296e-14 | 67.3367 | pfam13817 | DDE_Tnp_IS66_C | NA | cl16419 |
| pJBCL41_00334 | specific | 315644 | 48 | 117 | 3.50296e-13 | 64.2376 | pfam13007 | LZ_Tnp_IS66 | NA | cl15234 |
| pJBCL41_00335 | specific | 310376 | 5 | 104 | 2.34515e-47 | 146.826 | pfam05717 | TnpB_IS66 | NA | cl18171 |
| pJBCL41_00337 | superfamily | 320863 | 18 | 58 | 4.34679e-07 | 44.8271 | cl00259 | Sm_like superfamily | C | NA |
| pJBCL41_00339 | specific | 225116 | 18 | 257 | 4.14993e-86 | 261.654 | COG2206 | HDGYP | N | cl26425 |
| pJBCL41_00339 | superfamily | 331246 | 4 | 29 | 0.00238582 | 36.3579 | cl26425 | DUF3391 superfamily | N | NA |
| pJBCL41_00340 | specific | 314701 | 3 | 53 | 1.15477e-21 | 81.4262 | pfam11871 | DUF3391 | C | cl26425 |
| pJBCL41_00344 | specific | 316281 | 18 | 97 | 1.06146e-31 | 106.542 | pfam13744 | HTH_37 | NA | cl22854 |
| pJBCL41_00345 | superfamily | 328758 | 4 | 121 | 4.06058e-43 | 137.145 | cl21503 | ParE_toxin superfamily | NA | NA |
| pJBCL41_00346 | specific | 239737 | 13 | 141 | 8.87117e-56 | 173.049 | cd03768 | SR_ResInv | NA | cl02788 |
| pJBCL41_00346 | superfamily | 328727 | 153 | 190 | 4.6151e-07 | 44.6237 | cl21459 | HTH superfamily | NA | NA |
| pJBCL41_00347 | specific | 307598 | 596 | 988 | 4.36629e-127 | 390.704 | pfam01526 | DDE_Tnp_Tn3 | NA | cl14901 |
| pJBCL41_00347 | specific | 316241 | 10 | 173 | 3.08301e-38 | 139.996 | pfam13700 | DUF4158 | NA | cl18663 |
| pJBCL41_00349 | superfamily | 328722 | 3 | 36 | 0.00303965 | 31.9905 | cl21453 | PKc_like superfamily | NC | NA |
| pJBCL41_00350 | specific | 309066 | 630 | 877 | 8.84704e-45 | 162.256 | pfam03797 | Autotransporter | NA | cl22877 |
| pJBCL41_00351 | specific | 223999 | 130 | 253 | 0.00431239 | 37.7674 | COG1073 | FrsA | N | cl27027 |
| pJBCL41_00353 | superfamily | 331290 | 46 | 219 | 8.13833e-07 | 49.0535 | cl26469 | DsbD superfamily | N | NA |
| pJBCL41_00354 | specific | 315714 | 234 | 295 | 1.95813e-10 | 56.1843 | pfam13103 | TonB_2 | NA | cl10048 |
| pJBCL41_00354 | superfamily | 331285 | 93 | 185 | 4.85614e-05 | 44.9293 | cl26464 | Atrophin-1 superfamily | N | NA |
| pJBCL41_00354 | superfamily | 331068 | 16 | 214 | 0.00191276 | 39.7928 | cl26247 | DNA_pol3_delta2 superfamily | N | NA |
| pJBCL41_00355 | superfamily | 327377 | 2 | 206 | 1.77895e-50 | 175.062 | cl17041 | helicase_insert_domain superfamily | N | NA |
| pJBCL41_00356 | superfamily | 324572 | 160 | 232 | 9.23099e-07 | 45.3987 | cl10048 | TonB_C superfamily | NA | NA |
| pJBCL41_00357 | superfamily | 329065 | 105 | 187 | 6.5359e-08 | 49.6324 | cl23771 | Big_3_5 superfamily | NA | NA |
| pJBCL41_00358 | specific | 320690 | 78 | 263 | 6.28368e-76 | 229.001 | cd07331 | M48C_Oma1_like | NA | cl28898 |
| pJBCL41_00359 | superfamily | 304920 | 253 | 426 | 2.96198e-40 | 142.376 | cl23763 | MCP_signal superfamily | NA | NA |
| pJBCL41_00359 | specific | 312074 | 163 | 248 | 1.42084e-16 | 74.2966 | pfam08447 | PAS_3 | NA | cl25986 |
| pJBCL41_00359 | specific | 225112 | 31 | 255 | 5.44701e-16 | 76.8116 | COG2202 | PAS | NA | cl25988 |
| pJBCL41_00360 | specific | 185315 | 1 | 335 | 0 | 649.415 | PRK15417 | PRK15417 | NA | cl28330 |
| pJBCL41_00361 | specific | 306954 | 18 | 139 | 6.97981e-19 | 76.3946 | pfam00583 | Acetyltransf_1 | NA | cl17182 |
| pJBCL41_00362 | specific | 293861 | 42 | 259 | 6.40902e-150 | 417.34 | cd16303 | VIM_type_MBL-B1 | NA | cl23716 |
| pJBCL41_00363 | superfamily | 330977 | 1 | 172 | 2.65444e-138 | 382.31 | cl26156 | Acetyltransf_8 superfamily | NA | NA |
| pJBCL41_00364 | specific | 279265 | 3 | 94 | 1.75017e-28 | 98.8877 | pfam00893 | Multi_Drug_Res | NA | cl23754 |
| pJBCL41_00365 | specific | 184303 | 2 | 279 | 0 | 523.878 | PRK13753 | PRK13753 | NA | cl00219 |
| pJBCL41_00366 | specific | 306954 | 24 | 143 | 1.37001e-13 | 63.2978 | pfam00583 | Acetyltransf_1 | NA | cl17182 |
| pJBCL41_00368 | specific | 225853 | 1 | 213 | 2.78357e-69 | 212.384 | COG3316 | Rve | NA | cl26089 |
| pJBCL41_00369 | specific | 239737 | 23 | 147 | 5.70606e-54 | 168.042 | cd03768 | SR_ResInv | NA | cl02788 |
| pJBCL41_00369 | specific | 259851 | 159 | 200 | 4.66885e-13 | 60.8021 | cd00569 | HTH_Hin_like | NA | cl21459 |
| pJBCL41_00370 | specific | 307598 | 580 | 966 | 0 | 585.615 | pfam01526 | DDE_Tnp_Tn3 | NA | cl14901 |
| pJBCL41_00370 | specific | 316241 | 6 | 169 | 1.19181e-57 | 195.464 | pfam13700 | DUF4158 | NA | cl18663 |
| pJBCL41_00371 | specific | 274370 | 3 | 122 | 6.67628e-36 | 118.825 | TIGR02970 | succ_dehyd_cytB | NA | cl00881 |
| pJBCL41_00372 | specific | 307645 | 108 | 331 | 4.21438e-12 | 64.5404 | pfam01609 | DDE_Tnp_1 | NA | cl26088 |
| pJBCL41_00373 | superfamily | 294724 | 21 | 162 | 1.22572e-26 | 98.8025 | cl01162 | DUF417 superfamily | NA | NA |
| pJBCL41_00374 | specific | 225117 | 169 | 272 | 5.59162e-24 | 93.7769 | COG2207 | AraC | NA | cl26291 |
| pJBCL41_00374 | specific | 315518 | 2 | 147 | 1.32067e-16 | 75.4146 | pfam12852 | Cupin_6 | NA | cl21464 |
| pJBCL41_00375 | specific | 226950 | 11 | 278 | 1.41736e-16 | 79.8083 | COG4584 | COG4584 | NA | cl27787 |
| pJBCL41_00376 | specific | 307699 | 52 | 227 | 6.4385e-88 | 258.783 | pfam01695 | IstB_IS21 | NA | cl21455 |
| pJBCL41_00377 | specific | 234692 | 1 | 142 | 3.18217e-98 | 278.255 | PRK00222 | PRK00222 | NA | cl15841 |
| pJBCL41_00378 | specific | 237597 | 1 | 166 | 1.14351e-114 | 322.536 | PRK14054 | PRK14054 | NA | cl00366 |
| pJBCL41_00379 | superfamily | 330280 | 17 | 227 | 6.8725e-61 | 191.679 | cl25459 | GstA superfamily | NA | NA |
| pJBCL41_00380 | specific | 213364 | 32 | 322 | 3.07248e-141 | 400.361 | cd12830 | MtCorA-like | NA | cl00459 |
| pJBCL41_00382 | superfamily | 330398 | 2 | 337 | 0 | 565.904 | cl25577 | PKS_ER superfamily | NA | NA |
| pJBCL41_00383 | specific | 309158 | 30 | 112 | 5.97075e-24 | 89.0735 | pfam03928 | Haem_degrading | C | cl01249 |
| pJBCL41_00384 | specific | 311167 | 3 | 94 | 1.48566e-22 | 82.9805 | pfam07045 | DUF1330 | NA | cl22966 |
| pJBCL41_00386 | superfamily | 320948 | 26 | 140 | 0.000293448 | 37.9862 | cl00388 | Thioredoxin_like superfamily | NA | NA |
| pJBCL41_00387 | specific | 239027 | 18 | 197 | 6.0811e-90 | 261.025 | cd02109 | arch_bact_SO_family_Moco | NA | cl00199 |
| pJBCL41_00388 | specific | 99813 | 11 | 246 | 4.32018e-106 | 306.115 | cd06217 | FNR_iron_sulfur_binding_3 | NA | cl06868 |
| pJBCL41_00389 | specific | 274017 | 5 | 83 | 6.10501e-39 | 123.909 | TIGR02181 | GRX_bact | NA | cl00388 |
| pJBCL41_00390 | specific | 309397 | 2 | 89 | 1.25442e-33 | 110.996 | pfam04248 | NTP_transf_9 | NA | cl00998 |
| pJBCL41_00393 | specific | 274370 | 3 | 122 | 6.67628e-36 | 118.825 | TIGR02970 | succ_dehyd_cytB | NA | cl00881 |
| pJBCL41_00394 | specific | 281099 | 164 | 450 | 5.28757e-132 | 383.521 | pfam03050 | DDE_Tnp_IS66 | NA | cl24150 |
| pJBCL41_00394 | specific | 225970 | 81 | 240 | 2.13084e-25 | 101.755 | COG3436 | COG3436 | NA | cl26266 |
| pJBCL41_00394 | specific | 316346 | 456 | 493 | 4.82863e-17 | 74.2702 | pfam13817 | DDE_Tnp_IS66_C | NA | cl16419 |
| pJBCL41_00394 | specific | 315644 | 21 | 98 | 1.28099e-10 | 56.9188 | pfam13007 | LZ_Tnp_IS66 | NA | cl15234 |
| pJBCL41_00395 | specific | 310376 | 10 | 109 | 5.22711e-46 | 143.745 | pfam05717 | TnpB_IS66 | NA | cl18171 |
| pJBCL41_00396 | superfamily | 294724 | 21 | 162 | 1.22572e-26 | 98.8025 | cl01162 | DUF417 superfamily | NA | NA |
| pJBCL41_00397 | specific | 225117 | 169 | 272 | 5.59162e-24 | 93.7769 | COG2207 | AraC | NA | cl26291 |
| pJBCL41_00397 | specific | 315518 | 2 | 147 | 1.32067e-16 | 75.4146 | pfam12852 | Cupin_6 | NA | cl21464 |
| pJBCL41_00398 | superfamily | 333116 | 18 | 278 | 2.00252e-39 | 142.131 | cl28296 | NrfG superfamily | NA | NA |
| pJBCL41_00399 | specific | 309150 | 27 | 146 | 1.34467e-50 | 158.401 | pfam03918 | CcmH | NA | cl01179 |
| pJBCL41_00400 | specific | 239308 | 23 | 147 | 4.6039e-67 | 199.726 | cd03010 | TlpA_like_DsbE | NA | cl00388 |
| pJBCL41_00401 | specific | 226950 | 11 | 278 | 3.94349e-16 | 78.2675 | COG4584 | COG4584 | NA | cl27787 |
| pJBCL41_00402 | specific | 307699 | 52 | 227 | 6.4385e-88 | 258.783 | pfam01695 | IstB_IS21 | NA | cl21455 |
| pJBCL41_00404 | specific | 270307 | 28 | 238 | 5.46059e-84 | 251.377 | cd13589 | PBP2_polyamine_RpCGA009 | C | cl21456 |
| pJBCL41_00406 | specific | 307140 | 25 | 224 | 1.10017e-26 | 100.942 | pfam00857 | Isochorismatase | NA | cl00220 |
| pJBCL41_00407 | superfamily | 293530 | 82 | 186 | 3.90678e-28 | 101.528 | cl25201 | TetR_C_13 superfamily | NA | NA |
| pJBCL41_00407 | specific | 306858 | 13 | 59 | 2.80894e-07 | 45.1046 | pfam00440 | TetR_N | NA | cl27689 |
| pJBCL41_00408 | specific | 224715 | 3 | 193 | 9.70458e-42 | 141.364 | COG1802 | GntR | NA | cl28323 |
| pJBCL41_00409 | specific | 212491 | 8 | 239 | 9.14425e-63 | 195.965 | cd05233 | SDR_c | NA | cl25409 |
| pJBCL41_00410 | specific | 270307 | 23 | 291 | 2.97416e-90 | 271.022 | cd13589 | PBP2_polyamine_RpCGA009 | NA | cl21456 |
| pJBCL41_00411 | superfamily | 333220 | 12 | 349 | 1.10732e-83 | 273.439 | cl28400 | HemL superfamily | C | NA |
| pJBCL41_00412 | superfamily | 333220 | 22 | 435 | 3.80554e-159 | 475.283 | cl28400 | HemL superfamily | N | NA |
| pJBCL41_00413 | specific | 224045 | 4 | 247 | 8.64861e-68 | 209.727 | COG1120 | FepC | NA | cl28181 |
| pJBCL41_00414 | specific | 223682 | 30 | 164 | 3.7845e-31 | 114.284 | COG0609 | FepD | N | cl00454 |
| pJBCL41_00415 | specific | 307598 | 2 | 312 | 4.73951e-172 | 482.766 | pfam01526 | DDE_Tnp_Tn3 | N | cl14901 |
| pJBCL41_00417 | specific | 225076 | 4 | 151 | 8.69025e-11 | 58.8754 | COG2165 | PulG | NA | cl26823 |
| pJBCL41_00417 | superfamily | 309860 | 196 | 246 | 0.00124717 | 39.5943 | cl20035 | Shufflon_N superfamily | N | NA |
| pJBCL41_00418 | superfamily | 331290 | 14 | 553 | 4.57425e-82 | 267.462 | cl26469 | DsbD superfamily | NA | NA |
| pJBCL41_00420 | superfamily | 332058 | 40 | 268 | 1.77957e-14 | 71.1698 | cl27237 | Trypsin superfamily | NA | NA |
| pJBCL41_00424 | specific | 271180 | 174 | 351 | 9.78591e-57 | 182.884 | cd00799 | INT_Cre_C | NA | cl00213 |
| pJBCL41_00425 | superfamily | 330553 | 79 | 322 | 1.78954e-08 | 55.876 | cl25732 | SMC_N superfamily | NC | NA |
| pJBCL41_00425 | specific | 314581 | 9 | 113 | 6.21808e-08 | 49.9337 | pfam11740 | KfrA_N | NA | cl13226 |
| pJBCL41_00430 | superfamily | 333150 | 163 | 401 | 8.63239e-22 | 94.5965 | cl28330 | XerC superfamily | N | NA |
| pJBCL41_00431 | specific | 224900 | 18 | 284 | 2.75534e-44 | 150.63 | COG1989 | PulO | NA | cl26949 |
| pJBCL41_00432 | specific | 238088 | 5 | 120 | 3.26274e-24 | 89.1721 | cd00156 | REC | NA | cl19078 |
| pJBCL41_00433 | superfamily | 330553 | 95 | 144 | 0.00314435 | 36.5778 | cl25732 | SMC_N superfamily | NC | NA |
| pJBCL41_00437 | superfamily | 325143 | 86 | 174 | 0.000990284 | 40.3384 | cl11961 | ALDH-SF superfamily | NC | NA |
| pJBCL41_00438 | specific | 225581 | 62 | 319 | 4.38666e-39 | 137.718 | COG3039 | IS5 | NA | cl27014 |
| pJBCL41_00439 | superfamily | 331107 | 285 | 511 | 3.38344e-06 | 49.6316 | cl26286 | VirB4 superfamily | N | NA |
| pJBCL41_00441 | specific | 312026 | 33 | 145 | 1.28835e-31 | 112.181 | pfam08378 | NERD | NA | cl00516 |
| pJBCL41_00441 | superfamily | 332419 | 195 | 251 | 9.79416e-05 | 42.9227 | cl27598 | TOP1Bc superfamily | N | NA |
| pJBCL41_00442 | superfamily | 328784 | 26 | 74 | 0.00628374 | 33.1793 | cl21541 | OstA superfamily | C | NA |
| pJBCL41_00443 | superfamily | 330230 | 68 | 151 | 0.00237103 | 36.9285 | cl25409 | SDR superfamily | C | NA |
| pJBCL41_00444 | specific | 132992 | 3 | 96 | 2.72316e-22 | 91.5887 | cd06974 | TerD_like | N | cl18957 |
| pJBCL41_00444 | superfamily | 327490 | 118 | 200 | 0.0015749 | 39.0711 | cl18957 | TerD_like superfamily | C | NA |
| pJBCL41_00445 | superfamily | 328724 | 54 | 229 | 3.06411e-74 | 223.345 | cl21455 | P-loop_NTPase superfamily | NA | NA |
| pJBCL41_00445 | specific | 312099 | 24 | 51 | 2.68876e-10 | 53.9011 | pfam08483 | IstB_IS21_ATP | NA | cl26739 |
| pJBCL41_00446 | specific | 226950 | 34 | 289 | 5.30137e-39 | 142.981 | COG4584 | COG4584 | NA | cl27787 |
| pJBCL41_00446 | specific | 226950 | 229 | 499 | 5.99373e-15 | 74.8007 | COG4584 | COG4584 | NA | cl27787 |
| pJBCL41_00447 | specific | 308290 | 3 | 373 | 3.80028e-151 | 429.488 | pfam02595 | Gly_kinase | NA | cl00841 |
| pJBCL41_00451 | specific | 311934 | 150 | 202 | 4.70456e-09 | 51.088 | pfam08239 | SH3_3 | NA | cl17036 |
| pJBCL41_00455 | superfamily | 331747 | 2 | 97 | 1.4098e-14 | 71.0686 | cl26926 | LprI superfamily | C | NA |
| pJBCL41_00455 | superfamily | 330553 | 42 | 168 | 0.000340448 | 42.4702 | cl25732 | SMC_N superfamily | NC | NA |
| pJBCL41_00455 | specific | 320095 | 182 | 330 | 0.00478223 | 38.1538 | cd14964 | 7tm_GPCRs | N | cl28897 |
| pJBCL41_00456 | specific | 311452 | 4 | 169 | 2.66765e-68 | 205.257 | pfam07509 | DUF1523 | NA | cl06513 |
| pJBCL41_00457 | specific | 313378 | 222 | 421 | 2.77265e-93 | 279.144 | pfam10138 | vWA-TerF-like | NA | cl00057 |
| pJBCL41_00457 | specific | 132992 | 32 | 165 | 1.25895e-32 | 120.479 | cd06974 | TerD_like | NA | cl18957 |
| pJBCL41_00458 | specific | 316533 | 2 | 22 | 1.76946e-05 | 41.3469 | pfam14020 | DUF4236 | N | cl16543 |
| pJBCL41_00461 | specific | 308129 | 2 | 187 | 6.3796e-115 | 324.054 | pfam02342 | TerD | NA | cl18957 |
| pJBCL41_00462 | specific | 308129 | 2 | 187 | 1.54642e-123 | 346.011 | pfam02342 | TerD | NA | cl18957 |
| pJBCL41_00463 | superfamily | 324589 | 9 | 336 | 1.11486e-112 | 329.097 | cl10468 | TerC superfamily | NA | NA |
| pJBCL41_00464 | specific | 226316 | 11 | 149 | 6.03303e-36 | 120.967 | COG3793 | TerB | NA | cl11965 |
| pJBCL41_00465 | superfamily | 327490 | 199 | 396 | 3.69119e-93 | 277.895 | cl18957 | TerD_like superfamily | NA | NA |
| pJBCL41_00465 | superfamily | 327490 | 3 | 162 | 3.97485e-42 | 146.092 | cl18957 | TerD_like superfamily | NA | NA |
| pJBCL41_00466 | specific | 308129 | 2 | 191 | 8.62966e-65 | 197.324 | pfam02342 | TerD | NA | cl18957 |
| pJBCL41_00467 | superfamily | 328728 | 16 | 208 | 1.65754e-41 | 140.519 | cl21460 | HAD_like superfamily | NA | NA |
| pJBCL41_00468 | specific | 317930 | 3 | 297 | 3.82487e-122 | 352.624 | pfam15617 | C-C_Bond_Lyase | NA | cl21481 |
| pJBCL41_00469 | specific | 314199 | 14 | 255 | 8.01388e-128 | 366.448 | pfam11202 | PRTase_1 | NA | cl12751 |
| pJBCL41_00469 | specific | 317923 | 284 | 361 | 1.43494e-27 | 103.341 | pfam15608 | PELOTA_1 | NA | cl00600 |
| pJBCL41_00470 | superfamily | 328728 | 7 | 237 | 4.30093e-51 | 163.388 | cl21460 | HAD_like superfamily | NA | NA |
| pJBCL41_00471 | specific | 317924 | 36 | 221 | 3.23154e-79 | 241.281 | pfam15609 | PRTase_2 | NA | cl00309 |
| pJBCL41_00471 | superfamily | 315219 | 256 | 369 | 1.12819e-23 | 94.5451 | cl13881 | TRSP superfamily | NA | NA |
| pJBCL41_00472 | specific | 315930 | 190 | 319 | 3.79478e-25 | 98.9937 | pfam13365 | Trypsin_2 | NA | cl21584 |
| pJBCL41_00472 | specific | 132992 | 62 | 158 | 5.16664e-11 | 60.3876 | cd06974 | TerD_like | N | cl18957 |
| pJBCL41_00473 | specific | 317944 | 2 | 328 | 3.20708e-123 | 356.542 | pfam15632 | ATPgrasp_Ter | NA | cl25870 |
| pJBCL41_00474 | superfamily | 312721 | 13 | 53 | 0.00119527 | 35.1402 | cl07815 | DUF1974 superfamily | N | NA |
| pJBCL41_00480 | specific | 320084 | 79 | 120 | 3.55572e-19 | 73.9404 | cd16170 | MvaT_DBD | NA | cl28878 |
| pJBCL41_00482 | specific | 307613 | 145 | 398 | 8.2514e-14 | 70.1213 | pfam01555 | N6_N4_Mtase | NA | cl17173 |
| pJBCL41_00482 | specific | 316827 | 35 | 101 | 4.61213e-09 | 52.5586 | pfam14338 | Mrr_N | NA | cl26033 |
| pJBCL41_00486 | superfamily | 333132 | 129 | 313 | 1.00254e-08 | 59.0127 | cl28312 | UvrD superfamily | C | NA |
| pJBCL41_00486 | superfamily | 333132 | 516 | 675 | 0.000104291 | 45.5726 | cl28312 | UvrD superfamily | NC | NA |
| pJBCL41_00487 | superfamily | 333060 | 26 | 210 | 6.70716e-34 | 120.086 | cl28240 | Crp superfamily | NA | NA |
| pJBCL41_00489 | superfamily | 321924 | 35 | 190 | 1.85091e-25 | 96.2149 | cl02415 | DUF922 superfamily | NA | NA |
| pJBCL41_00490 | superfamily | 331071 | 355 | 515 | 0.00317405 | 39.3512 | cl26250 | TPR_6 superfamily | N | NA |
| pJBCL41_00491 | superfamily | 325063 | 58 | 310 | 5.13566e-07 | 50.3241 | cl11502 | Ter superfamily | NA | NA |
| pJBCL41_00492 | specific | 223861 | 23 | 147 | 8.63537e-09 | 53.1243 | COG0790 | TPR | NC | cl27881 |
| pJBCL41_00494 | specific | 311281 | 37 | 109 | 7.29718e-05 | 38.6361 | pfam07238 | PilZ | N | cl01260 |
| pJBCL41_00497 | superfamily | 320896 | 29 | 125 | 1.13521e-08 | 49.0092 | cl00314 | Ribosomal_S10 superfamily | NA | NA |
| pJBCL41_00498 | specific | 223736 | 61 | 224 | 1.06332e-08 | 53.2967 | COG0664 | Crp | NA | cl28240 |
| pJBCL41_00501 | superfamily | 333132 | 32 | 1115 | 8.57251e-73 | 263.554 | cl28312 | UvrD superfamily | NA | NA |
| pJBCL41_00502 | superfamily | 333072 | 13 | 897 | 1.60959e-138 | 434.085 | cl28252 | COG3893 superfamily | NA | NA |
| pJBCL41_00503 | specific | 225110 | 22 | 246 | 2.34173e-43 | 147.051 | COG2200 | EAL | NA | cl27669 |
| pJBCL41_00504 | superfamily | 328724 | 68 | 325 | 1.17734e-38 | 140.478 | cl21455 | P-loop_NTPase superfamily | N | NA |
| pJBCL41_00505 | superfamily | 332997 | 15 | 96 | 3.35097e-05 | 41.1888 | cl28177 | SkfB superfamily | N | NA |
| pJBCL41_00506 | specific | 226950 | 34 | 289 | 5.30137e-39 | 142.981 | COG4584 | COG4584 | NA | cl27787 |
| pJBCL41_00506 | specific | 226950 | 229 | 499 | 5.99373e-15 | 74.8007 | COG4584 | COG4584 | NA | cl27787 |
| pJBCL41_00507 | superfamily | 328724 | 54 | 229 | 3.06411e-74 | 223.345 | cl21455 | P-loop_NTPase superfamily | NA | NA |
| pJBCL41_00507 | specific | 312099 | 24 | 51 | 2.68876e-10 | 53.9011 | pfam08483 | IstB_IS21_ATP | NA | cl26739 |
| pJBCL41_00508 | superfamily | 332997 | 24 | 296 | 3.93641e-10 | 59.7654 | cl28177 | SkfB superfamily | C | NA |
| pJBCL41_00510 | superfamily | 332997 | 26 | 386 | 4.5364e-13 | 69.7806 | cl28177 | SkfB superfamily | NA | NA |
| pJBCL41_00512 | specific | 316188 | 90 | 192 | 2.29249e-06 | 44.2962 | pfam13640 | 2OG-FeII_Oxy_3 | NA | cl21496 |
| pJBCL41_00513 | superfamily | 332997 | 14 | 355 | 9.8364e-11 | 62.4618 | cl28177 | SkfB superfamily | NA | NA |
| pJBCL41_00515 | specific | 226950 | 11 | 278 | 1.41736e-16 | 79.8083 | COG4584 | COG4584 | NA | cl27787 |
| pJBCL41_00516 | specific | 307699 | 52 | 227 | 6.4385e-88 | 258.783 | pfam01695 | IstB_IS21 | NA | cl21455 |
| pJBCL41_00517 | superfamily | 332993 | 18 | 77 | 0.000190243 | 40.5252 | cl28173 | PRK13758 superfamily | NC | NA |
| pJBCL41_00518 | superfamily | 332403 | 1054 | 1121 | 0.00252159 | 41.8349 | cl27582 | PRK05865 superfamily | N | NA |
| pJBCL41_00519 | superfamily | 332993 | 11 | 83 | 8.77812e-07 | 51.2465 | cl28173 | PRK13758 superfamily | C | NA |
| pJBCL41_00519 | superfamily | 332997 | 23 | 149 | 1.21988e-05 | 47.8242 | cl28177 | SkfB superfamily | C | NA |
| pJBCL41_00520 | superfamily | 332997 | 20 | 370 | 8.23579e-09 | 57.4542 | cl28177 | SkfB superfamily | NA | NA |
| pJBCL41_00521 | superfamily | 332997 | 14 | 326 | 2.60563e-09 | 57.8394 | cl28177 | SkfB superfamily | NA | NA |
| pJBCL41_00524 | specific | 223714 | 20 | 199 | 1.70456e-21 | 94.4333 | COG0641 | AslB | C | cl28177 |
| pJBCL41_00525 | specific | 223714 | 3 | 348 | 2.70287e-26 | 107.915 | COG0641 | AslB | NA | cl28177 |
| pJBCL41_00526 | specific | 223714 | 1 | 386 | 7.8509e-38 | 140.272 | COG0641 | AslB | NA | cl28177 |
| pJBCL41_00527 | superfamily | 331644 | 14 | 73 | 1.08095e-07 | 50.1172 | cl26823 | N_methyl superfamily | C | NA |
| pJBCL41_00528 | specific | 225076 | 1 | 144 | 3.34765e-11 | 58.105 | COG2165 | PulG | NA | cl26823 |
| pJBCL41_00529 | superfamily | 333156 | 18 | 387 | 4.00081e-07 | 51.4431 | cl28336 | PulF superfamily | NA | NA |
| pJBCL41_00532 | specific | 225363 | 113 | 560 | 4.59844e-113 | 346.625 | COG2804 | PulE | NA | cl28318 |
| pJBCL41_00536 | superfamily | 332547 | 267 | 541 | 2.07139e-29 | 117.483 | cl27726 | Secretin superfamily | NA | NA |
| pJBCL41_00538 | superfamily | 328734 | 207 | 298 | 0.00362638 | 38.4093 | cl21469 | HDc superfamily | C | NA |
| pJBCL41_00540 | superfamily | 331107 | 47 | 93 | 0.000721376 | 41.9276 | cl26286 | VirB4 superfamily | NC | NA |
| pJBCL41_00541 | superfamily | 327527 | 73 | 122 | 0.000434658 | 39.5072 | cl19237 | DUF45 superfamily | NC | NA |
| pJBCL41_00543 | superfamily | 328724 | 54 | 229 | 3.06411e-74 | 223.345 | cl21455 | P-loop_NTPase superfamily | NA | NA |
| pJBCL41_00543 | specific | 312099 | 24 | 51 | 2.68876e-10 | 53.9011 | pfam08483 | IstB_IS21_ATP | NA | cl26739 |
| pJBCL41_00544 | specific | 226950 | 34 | 289 | 5.30137e-39 | 142.981 | COG4584 | COG4584 | NA | cl27787 |
| pJBCL41_00544 | specific | 226950 | 229 | 499 | 5.99373e-15 | 74.8007 | COG4584 | COG4584 | NA | cl27787 |
| pJBCL41_00549 | superfamily | 314319 | 20 | 301 | 3.80658e-14 | 71.5266 | cl20092 | DUF3150 superfamily | NA | NA |
| pJBCL41_00549 | superfamily | 330571 | 326 | 377 | 0.0019745 | 39.6664 | cl25750 | DamX superfamily | NC | NA |
| pJBCL41_00552 | superfamily | 320736 | 550 | 639 | 4.26144e-08 | 53.0983 | cl00057 | vWFA superfamily | C | NA |
| pJBCL41_00556 | superfamily | 331165 | 35 | 366 | 5.01805e-21 | 92.7751 | cl26344 | CobS_N superfamily | NA | NA |
| pJBCL41_00563 | superfamily | 321087 | 22 | 294 | 2.45108e-09 | 56.795 | cl00641 | Cas4_I-A_I-B_I-C_I-D_II-B superfamily | NA | NA |
| pJBCL41_00565 | superfamily | 226990 | 58 | 318 | 5.16502e-05 | 44.8353 | cl26703 | COG4643 superfamily | C | NA |
| pJBCL41_00566 | specific | 275209 | 62 | 406 | 1.0663e-176 | 498.906 | TIGR04416 | group_II_RT_mat | NA | cl26764 |
| pJBCL41_00567 | superfamily | 331543 | 23 | 207 | 4.2052e-26 | 103.613 | cl26722 | Spo0J superfamily | NA | NA |
| pJBCL41_00568 | specific | 224113 | 12 | 244 | 1.16454e-37 | 133.799 | COG1192 | BcsQ | NA | cl27521 |
| pJBCL41_00569 | superfamily | 331332 | 23 | 342 | 7.85271e-07 | 52.0655 | cl26511 | Neuromodulin_N superfamily | NC | NA |
| pJBCL41_00570 | superfamily | 331543 | 22 | 223 | 1.45121e-46 | 156.519 | cl26722 | Spo0J superfamily | C | NA |
| pJBCL41_00572 | specific | 306560 | 7 | 118 | 2.67505e-24 | 89.1328 | pfam00072 | Response_reg | NA | cl19078 |
| pJBCL41_00573 | specific | 279866 | 21 | 157 | 3.49375e-11 | 57.2248 | pfam01584 | CheW | NA | cl00256 |
| pJBCL41_00574 | specific | 223910 | 336 | 661 | 1.00954e-12 | 70.4048 | COG0840 | Tar | N | cl28165 |
| pJBCL41_00575 | superfamily | 332069 | 13 | 250 | 2.02351e-44 | 150.514 | cl27248 | CheR_N superfamily | NA | NA |
| pJBCL41_00576 | superfamily | 332149 | 959 | 1814 | 1.8687e-96 | 328.906 | cl27328 | H-kinase_dim superfamily | NA | NA |
| pJBCL41_00576 | specific | 238088 | 1839 | 1951 | 1.7806e-29 | 113.825 | cd00156 | REC | NA | cl19078 |
| pJBCL41_00577 | specific | 307485 | 127 | 302 | 6.76379e-31 | 114.056 | pfam01339 | CheB_methylest | NA | cl03170 |
| pJBCL41_00584 | superfamily | 331967 | 330 | 520 | 3.40324e-27 | 111.183 | cl27146 | Sigma70_r3 superfamily | N | NA |
| pJBCL41_00585 | specific | 238082 | 1 | 368 | 1.91822e-79 | 247.424 | cd00140 | beta_clamp | NA | cl27370 |
| pJBCL41_00586 | superfamily | 321524 | 102 | 200 | 0.00697273 | 36.4955 | cl01482 | CpxP_like superfamily | N | NA |
| pJBCL41_00587 | specific | 226950 | 34 | 289 | 5.30137e-39 | 142.981 | COG4584 | COG4584 | NA | cl27787 |
| pJBCL41_00587 | specific | 226950 | 229 | 499 | 5.99373e-15 | 74.8007 | COG4584 | COG4584 | NA | cl27787 |
| pJBCL41_00588 | superfamily | 328724 | 54 | 229 | 3.06411e-74 | 223.345 | cl21455 | P-loop_NTPase superfamily | NA | NA |
| pJBCL41_00588 | specific | 312099 | 24 | 51 | 2.68876e-10 | 53.9011 | pfam08483 | IstB_IS21_ATP | NA | cl26739 |
| pJBCL41_00589 | superfamily | 328777 | 34 | 109 | 0.000780396 | 38.6686 | cl21532 | NADAR superfamily | NC | NA |
| pJBCL41_00590 | superfamily | 331580 | 69 | 431 | 2.42755e-21 | 95.938 | cl26759 | Exonuc_X-T_C superfamily | NA | NA |
| pJBCL41_00591 | superfamily | 304554 | 14 | 243 | 1.63186e-82 | 246.896 | cl22428 | E1_enzyme_family superfamily | NA | NA |
| pJBCL41_00592 | superfamily | 326315 | 6 | 227 | 2.23622e-43 | 145.608 | cl14019 | Prok-E2_D superfamily | NA | NA |
| pJBCL41_00594 | superfamily | 326316 | 5 | 62 | 3.56126e-18 | 69.9593 | cl14020 | Prok_Ub superfamily | NA | NA |
| pJBCL41_00595 | superfamily | 274758 | 4 | 109 | 1.01739e-06 | 44.6827 | cl14021 | PRTRC_E superfamily | NA | NA |
| pJBCL41_00596 | superfamily | 331543 | 34 | 593 | 5.96008e-117 | 358.639 | cl26722 | Spo0J superfamily | NA | NA |
| pJBCL41_00600 | superfamily | 321363 | 1 | 112 | 0.00953034 | 35.4752 | cl01135 | ABC_trans_aux superfamily | C | NA |

NA stands for no information available.
