## Supplemental Table 2 for "Megaplasmids on the Rise: Combining Sequencing Approaches to Fully Resolve a Carbapenemase-Encoding Plasmid in a Proposed Novel *Pseudomonas* Species"

**Table S2**. Matrix of the number of orthogroups shared by each megaplasmid’s protein sequences.

|  | AR439_plasmid | P19E3_p1 | RW109_plasmid | XWY-1_plasmid | pBM413 | pJB37 | pJBCL41 | pOZ176 | pQBR103 | pSY153-MDR |
| --- | --- | --- | --- | --- | --- | --- | --- | --- | --- | --- |
| AR439_plasmid |  | 466 | 451 | 44 | 456 | 466 | 276 | 467 | 242 | 452 |
| P19E3_p1 | 466 |  | 437 | 49 | 461 | 465 | 267 | 453 | 244 | 455 |
| RW109_plasmid | 451 | 437 |  | 48 | 425 | 440 | 247 | 453 | 218 | 434 |
| XWY-1_plasmid | 44 | 49 | 48 |  | 46 | 56 | 63 | 56 | 33 | 46 |
| pBM413 | 456 | 461 | 425 | 46 |  | 469 | 273 | 460 | 236 | 494 |
| pJB37 | 466 | 465 | 440 | 56 | 469 |  | 274 | 507 | 243 | 461 |
| pJBCL41 | 276 | 267 | 247 | 63 | 273 | 274 |  | 277 | 335 | 275 |
| pOZ176 | 467 | 453 | 453 | 56 | 460 | 507 | 277 |  | 239 | 460 |
| pQBR103 | 242 | 244 | 218 | 33 | 236 | 243 | 335 | 239 |  | 236 |
| pSY153-MDR | 452 | 455 | 434 | 46 | 494 | 461 | 275 | 460 | 236 |  |
